# Electrochemotherapy with bleomycin associated with doxorubicin induces tumor regression and decreases the proliferative index in canine cutaneous squamous cell carcinomas

**DOI:** 10.1101/474262

**Authors:** Denner S. Anjos, Cynthia Bueno, Larissa F. Magalhães, Georgia M. Magalhães, Ewaldo Mattos-Junior, Marcela M.R. Pinto, Andrigo B. De Nardi, Carlos H.M. Brunner, Antonio F. Leis-Filho, Carlos E. Fonseca-Alves, Sabryna G. Calazans

**Affiliations:** Veterinary Science Graduate Program, University of Franca (UNIFRAN), Franca, Brazil; Department of Veterinary Clinic and Surgery, São Paulo State University (UNESP), Jaboticabal, Brazil; Department of Veterinary Pathology, University of Franca (UNIFRAN), Franca, Brazil; Federal Institute of Education, Science and Technology of the South of Minas Gerais - Muzambinho, Minas Gerais, Brazil; Pathologist veterinarian at the CEVEPAT Laboratory, Botucatu, SP; Department of Veterinary Clinic of University Paulista, São Paulo, Brazil; Department of Veterinary Clinic, School of Veterinary Medicine and Animal Science, São Paulo State University (UNESP), Botucatu, São Paulo, Brazil

**Keywords:** Apoptotic markers, cutaneous carcinoma, electroporation, proliferative index, tumor size

## Abstract

Canine cutaneous squamous cell carcinoma (cSCC) is the most common skin cancer in dogs, and due to its low metastatic rate, local treatments such as electrochemotherapy (ECT) promote disease control or even complete remission and increase the survival time in most cases. This study aimed to evaluate the expression of BAX, Bcl-2, and Ki67 and clinical parameters in dogs with cSCC subjected to ECT. A prospective clinical nonrandomized study was performed in dogs with naturally occurring cSCC treated with ECT. Eighteen lesions (from 11 dogs) were selected, independent of breed, sex and age. The ECT protocol consisted of bleomycin plus doxorubicin followed by electric pulses characterized by 8 biphasic electric pulses lasting 100 ms, 1 Hz and 1000 V/cm. Among the 18 lesions, the lesion volume significantly decreased after treatment (p=0.04). The tumor size at D0 had no impact on survival time or prognosis (P>0.05). A decreased mitotic index was observed at compared with D0 (P=0.019). We also observed more intratumoral necrosis at D21 compared to D0 (P=0.041). The median expression level of Ki67 was 277.96 at D0 and 193.92 at D21. Thus, tumor samples had a lower proliferative index after ECT (D21) (P=0.031). The survival times of subjects with Ki67 values lower and higher than the Ki67 median value were not significantly different (P>0.05). Regarding apoptotic markers, there was no significant difference in BAX or Bcl-2 expression between D0 and D21 (P>0.05) or in overall survival between subjects with different levels of apoptotic markers. Furthermore, a positive correlation was observed between BAX and Bcl-2 before ECT (D0) (P=0.0379, r=0.5067). In conclusion, there was no change in BAX and Bcl-2 protein expression levels in response to ECT at the time points evaluated, and ECT was able to reduce tumor volume and cellular proliferation in cSCC.

## Introduction

Canine cutaneous squamous cell carcinoma (cSCC) is the most common skin cancer in dogs, and the most important etiological factor is chronic sunlight exposure. On the other hand, canine papillomavirus infection, chronic inflammation and immunosuppression might be involved in the development of cSCC [1–4]. cSCC is the second most common cutaneous tumor in dogs [5], with an incidence between 3-20% [6,7]; however, the incidence of cSCC depends on the geographic area [5]. In humans and dogs, actinic keratosis is an SCC precursor lesion with more than 80% of the SCC cases in humans being derived from previous actinic keratosis [8, 9].

Generally, cSCCs are locally invasive tumors with a low metastatic rate (13%) [10–13], with is similar to the behavior of SCCs humans, in whom the metastatic rate is 5%; metastasis mainly occurs in locoregional lymph nodes [14–16]. Due to this low risk of metastasis, local treatments such as surgery, cryotherapy, radiotherapy, photodynamic therapy and electrochemotherapy (ECT) promote disease control and increase the survival time in most cases [1,17].

ECT is able to induce an inflammatory response, necrosis, scar tissue and apoptosis in tumors [18–20]. This process is a genetically programmed process for the elimination of damaged cells, and its onset is controlled by numerous interrelating processes that are influenced by extrinsic and intrinsic signals that converge in an effector pathway. Alterations in these pathways are an important in the process of tumorigenesis, leading to the persistence of neoplastic cells and the promotion of progression and metastasis [21,22].

BAX (Bcl-2 associated X protein) and Bcl-2 are important proteins in the BCL-2 family, which consists of pro- and anti-apoptotic proteins. The BAX/Bcl-2 cross-regulation controls apoptosis, cell survival or cellular proliferation [23,24]. Several studies have determined the expression level of a single apoptosis-associated protein, such as BAX and/or BCL-2, by immunohistochemistry or flow cytometry and have correlated its expression with the prognosis of mammary tumors and lymphoma [25,26]. However, conflicting data have been observed among the studies [25–27]. In feline cutaneous SCCs, neither BAX nor BCL-2 expression was detected. However, in basal cell tumors, BCL-2 expression was higher (23/24) than that of BAX, which was expressed only in seven out of 24 tumors. For the tumors that expressed both BAX and BCL-2, the BAX:BCL-2 ratio was low [28].

Another important factor to be evaluated in cSCC is cellular proliferation, which is measured by the level of Ki67. It is a nuclear protein that is used to detect proliferative cells because it is present in proliferating cells during the late G1, S, G2 and M phases of the cell cycle, and it is correlated with poor prognosis in dogs [29–32]. It is known that the high expression level of Ki67 is correlated with a poorer prognosis in dogs with several tumor types as well as in humans, such as mammary tumors, mast cell tumors, perianal tumors, oral and cSCCs [29–37].

ECT has gained popularity in recent years in veterinary medicine as well as in human medicine [38,39]. ECT is a combination of chemotherapy and the localized delivery of electric pulses to the tumor nodule. ECT aims to increase antineoplastic drug diffusion into the cell after cell membrane electroporation, thus increasing its cytotoxicity [17].

To the best of the authors’ knowledge, this is the first study to evaluate the expression of BAX, Bcl-2, and Ki67 in dogs with cSCC that underwent ECT. This study aimed to evaluate the clinical parameters, proliferative index, and expression levels of BAX and Bcl-2 in dogs with cSCC that underwent ECT.

## Materials and Methods

### Ethical approval

This study was performed in accordance with the National and International Recommendations for the Care and Use of Animals. All procedures were performed after receiving the approval of the Institutional Animal Ethics Committee (CEUA/UNIFRAN, #033/15).

### Study design

A prospective clinical nonrandomized study was performed in dogs that presented to the Veterinary Teaching Hospital of Sao Paulo State University (UNESP) and the Veterinary Teaching Hospital of University of Franca (UNIFRAN) with naturally occurring cSCC and that were treated with ECT. Consent was obtained from the dogs’ owners to perform the treatment. The recommendations made by Campana et al. [40] for reporting clinical studies on ECT were followed.

### Inclusion criteria and animal selection

All dogs enrolled in this study fulfilled the following characteristics: a histopathologically confirmed diagnosis of stage T1 cSCC (according to the World Health Organization) [41]; the absence of distant metastases; the compliance of the owner with follow-up after 21 days; the owner’s permission to perform biopsies before (D0) and after ECT (D21).

Eighteen lesions (from 11 dogs) were selected, independent of breed, sex and age. The eligibility inclusion criteria were as follows: dogs with clinically staged cSCC and complete physical examinations, laboratory exams, and fine needle aspiration of regional lymph nodes. Three-view thoracic radiography and abdominal ultrasonography were also performed.

### Electrochemotherapy protocol

Bleomycin (Cinaleo®-Meizler, Barueri-SP) was diluted in 5 mL of saline solution and administered intravenously (IV) at a dose of 15,000 UI/m^2^. Five minutes after the administration of the bleomycin, doxorubicin (Cloridrato de Doxorrubicina^®^ - Eurofarma Laboratórios S.A. Ribeirao Preto – SP) was diluted in 25 mL of saline solution and administered intravenously at a dose of 30 mg/m², followed by sequences of 8 biphasic electric pulses lasting 100 ms each, with a frequency of 1 Hz, 1000 V, generated by an LC BK-100 portable electroporator and delivered by six needle electrodes with 0.3 mm distance between them; the needle electrodes were arranged in rows (parallel array) until they achieved complete coverage of the tumor. The procedure lasted until 28 minutes, according to a previous report [42,43].

All ECT procedures were administered under general anesthesia using propofol (5 mg/kg) IV followed by endotracheal intubation; anesthesia was maintained with isoflurane. All animals received postoperative analgesia, including the IV injection of meloxicam (0.2 mg/kg) and tramadol (2 mg/kg).

### Tumor response

The total volume of the neoplasm was calculated by the following formula: *π x length x width x height/6;* tumor volume was measured at D0 and D21 using a digital pachymeter [29,30]. The same evaluator measured tumor volume to exclude measurement bias. Complete remission (CR) was defined as a total reduction in measured tumor volume, while partial remission (PR) was defined as ≥ 30% reduction in tumor volume. Progressive disease (PD) was defined as ≥ 20% increase in tumor volume or new lesions, and stable disease (SD) was defined as ˂ 30% reduction in tumor volume or ˂ 20% increase in tumor volume [43]. CR and RP were considered “favorable” responses, while PD and SD were considered “unfavorable” responses.

### Histopathological evaluation

The first sample was collected immediately before the first session of ECT (D0). The second sample was obtained on day 21 (D21) after ECT. A 6-mm punch biopsy was used to obtain the tumor samples. All samples were immediately placed in 10% formalin for 24 hours, followed by 70% alcohol until paraffinized sections were prepared. All tumor samples were classified according to Gross et al. [5], as follows: 1) well-differentiated SCC, presenting centralized accumulatio of compact laminated keratin or keratin pearls, with keratinization progressing through the granular cell layer as in the normal epidermis and the keratinized centers of the lobules undergoing necrosis and becoming infiltrated by neutrophils or 2) poorly differentiated SCC presenting smaller epithelial structures, cords and nests rather than large islands of squamous epithelial cells, moderate to high mitotic activity, and no keratin pearls.

### Immunohistochemistry

Immunohistochemical staining for BAX, BCL-2 and Ki67 antibodies was performed in all cSCCs in the original biopsy samples (D0) and in the biopsy specimens collected 21 days after the first ECT (D21). Immunohistochemical staining was performed using the peroxidase method and 3,3’ diaminobenzidine tetrachloride (DAB). The slides were dewaxed in xylol and rehydrated in graded ethanol. For antigen retrieval, the slides were incubated in citrate buffer (pH 6.0) in a pressure cooker (Pascal, Agilent Technologies, Santa Clara, CA, USA). The endogenous peroxidase was blocked with a commercial solution (Protein Block, Agilent Technologies, Santa Clara, CA, USA), and the samples were incubated with primary antibodies overnight at 4ºC. Anti-BAX (mouse monoclonal, Santa Cruz Biotechnology, Dallas, TX, USA) was detected with a monoclonal antibody at a 1:200 dilution overnight. Bcl-2 (mouse monoclonal, Santa Cruz Biotechnology, Dallas, TX, USA) was a monoclonal mouse antibody used at a 1:400 dilution overnight, and anti-Ki67 (MIB-1, Agilent Technologies, Santa Clara, CA, USA) with a monoclonal mouse antibody used at a 1:50 dilution, applied overnight. After incubation with the above antibodies, the slides were placed on an autostainer platform (Agilent Technologies, Santa Clara, CA, USA). Then, the sections were counterstained with Harris’s hematoxylin, dehydrated, and mounted. The anti-Ki67 antibody cross-reactivity with canine tissue was provided by the manufacturer in the antibody datasheet. For the anti-Bcl-2 and anti-BAX antibodies, we performed western blotting to show the antibody reactivity.

Negative controls were run for all antibodies with a mouse universal negative control (Dako, Carpinteria, CA, USA) according to the manufacturer’s instructions. The positive control consisted of normal lymph nodes for all antibodies according to Protein Atlas guidelines (www.proteinatlas.org).

### Immunohistochemical evaluation

The immunohistochemical slides were examined with a light microscope using an ocular grid 26-mm in diameter (Leica Microscopy DMLB, HC, PLAN 10x/20, 4”x5”) at 400X magnification. The immunoexpression levels of BAX, Bcl-2 and Ki67 were established by counting the number of stained cells and considering the number of positive cells in relation to the total number of cells inside the ocular grid per five random high-power fields (400x). The samples were classified as 0 (absence of stained cells), 1 (< 10% stained cells), 2 (10-25% stained cells), 3 (26-50% stained cells) and 4 (> 50% stained cells) according to Fonseca-Alves et al. [44]. For the evaluation of the scores, cytoplasmic staining for BAX and Bcl-2 and nuclear staining for Ki67 were considered.

### Western blotting

The western blot analysis was performed to validate the protein cross-reactivity with canine tissue. The procedures were performed according to a previously published method [45]. Briefly, two cSCC tissue biopsies were collected and homogenized (Polytron homogenizer, Kinematica, Lucerne, Switzerland) for 30 seconds at 4°C in RIPA Buffer (Millipore, Burlington, MA, USA). Total protein was extracted and quantified as described by Bradford [46]. A total of 70 μg of protein was subjected to electrophoresis and then transferred to nitrocellulose membranes (Sigma Chemical Co., St. Louis, MO). The blots were blocked with 3% bovine serum albumin in TBS-T for 2 hours and incubated overnight with BAX (mouse monoclonal, Santa Cruz Biotechnology, Dallas, TX, USA) and Bcl-2 (mouse monoclonal, Santa Cruz Biotechnology, Dallas, TX, USA) antibodies. Goat anti-β-actin antibody (1:1,000; sc-1615, Santa Cruz Biotechnology, Santa Cruz, CA, USA) was used as a control. A horseradish peroxidase-conjugated secondary antibody was incubated with the blots for 1 hour, and the blots were visualized via chemiluminescence (Amersham ECL Select Western Blotting Detection Reagent, GE Healthcare).

### Statistical analysis

For statistical purposes, we calculated the median value of each clinical, histopathological and immunohistochemical parameter and classified the value as “low” when it was lower than median and “high” when the value was greater than the median. Then, we compared the survival between dogs with low and high expression values of each parameter through the Kaplan-Myer method. To evaluate the association between the tumor stage (≤T2 x

>T2) before treatment and after ECT, Fisher’s exact test was applied. Furthermore, Spearman’s correlations between the expression levels of BAX, Bcl-2 and Ki67 were performed to compare the levels at two time points, namely, before and after ECT. Commercial software (GraphPad Prism^®^ 6.0 – GraphPad Software, Inc. 2015) was used for the statistical analysis. P values <0.05 were considered significant.

## Results

### Clinical data

Mixed breed dogs (n=5) were the dogs most commonly affected by cSCC, followed by American Pit Bulls (n=3), Boxers (n=2) and English Pointers (n=1). All subjects had sparse fur and lightly pigmented skin. The mean age of the animals was 7.5 (± 2.29) years, with females being the more common sex (n=10). Five animals had more than one lesion. Thus, we investigated 18 lesions in 11 subjects.

The most commonly affected sites observed were the abdominal region (n=10), followed by the thoracic region (n=5), axillar region (n=1), preputial region (n=1) and tibial region (n=1). Among the 11 subjects, only three had regional lymph node involvement at the time of diagnosis. There was no difference in survival time between subjects with and without lymph node involvement (S Fig 1A) (P> 0.05).

**Fig. 1.**
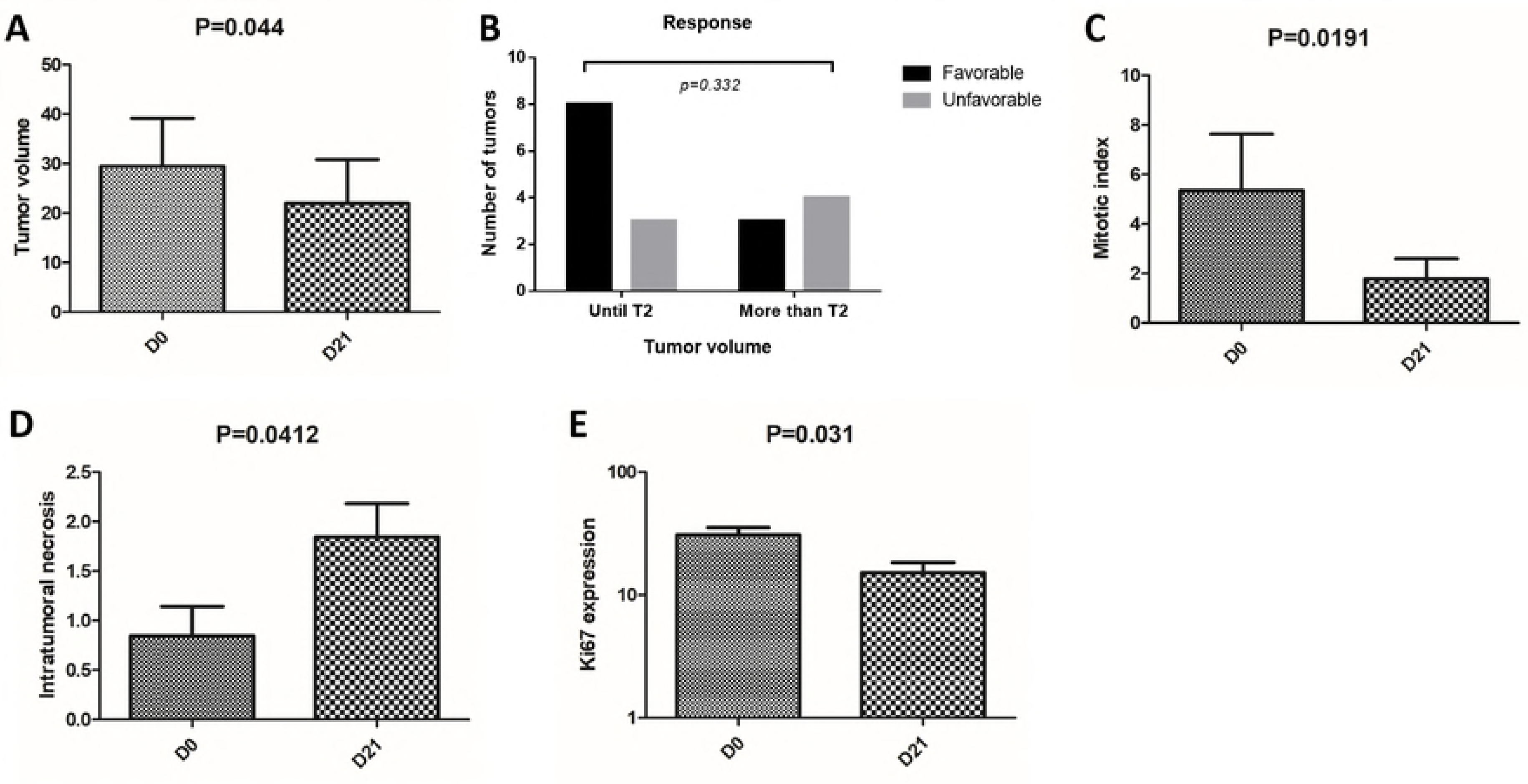
Evaluation of tumor volume, clinical response according to tumor volume, mitotic index, intratumoral necrosis and Ki67 expression before and after treatment with ECT in dogs with cSCC. (A) The volume before the treatment ranged from 0.14 to 112.9 cm³ (median of 4.64), and the volume significantly decreased after treatment (p=0.04), ranging from 0.11 to 118.2 cm³ (median of 1.49). (B) Evaluation of tumor size at D0 as prognostic factor in dogs with tumors smaller and larger than 5 cm^3^. The volume at D0 had no impact on survival time and had no prognostic value (P>0.05), and the response to ECT was not significant (p=0.332). (C) A decreased mitotic index at D21 (median 1.5, ranging 0 to 20) (P=0.019) was observed. (D) Greater intratumoral necrosis was observed in tumor samples at D21 compared to those at D0 (P=0.041). (E) Lower proliferative index (p=0.031) was observed at D21 when compared to D0.

Among the 18 lesions, the volume before the treatment ranged from 0.14 to 112.9 cm³ (median of 4.64), and the volume decreased significantly after treatment (p=0.04), ranging from 0.11 to 118.2 cm³ (median of 1.49) (Table 1) (Fig 1A).

**Table 1.** Records from the subjects that underwent ECT regarding the clinical data, tumor localization, stage and tumor size.

| ID | Breed | Age | Sex | Number of tumors | Tumor localization | Histopathological evaluation | TNM | Tumor size at D0 (cm<sup>3</sup>) | Tumor size at D21 (cm<sup>3</sup>) | Outcome | Number of sessions |
| --- | --- | --- | --- | --- | --- | --- | --- | --- | --- | --- | --- |
| 1 | Pitbull | 8 | M | 2 (a, b) | Axillar (a)<br>Prepucial (b) | poorly differentiated<br>well differentiated | T2N0M0<br>T2N0M0 | 3.75<br>19.23 | 0.91<br>6.44 | PR<br>PR | 3 / S |
| 2 | Pitbull | 9 | F | 1 | Abdominal | well differentiated | T2N0M0 | 5.04 | 0.73 | PR | 2 |
| 3 | Boxer | 2 | F | 1 | Thorax lateral | well differentiated | T3N0M0 | 68.49 | 65.94 | SD | 2 / S |
| 4 | Mixed | 10 | F | 2 (a, b) | Thorax lateral (a)<br>Abdominal (b) | poorly differentiated<br>well differentiated | T2N0M0<br>T3N0M0 | 2.3<br>14.15 | 0.278<br>1.68 | PR<br>PR | 1 / S |
| 5 | Mixed | 7 | F | 1 | Abdominal | well differentiated | T3N0M0 | 93.57 | 118.24 | PD | 2 |
| 6 | Pitbull | 9 | F | 4 (a, b, c, d) | Thorax lateral (a)<br>Abdominal (b)<br>Abdominal (c)<br>Abdominal (d) | well differentiated<br>poorly differentiated<br>well differentiated<br>well differentiated | T1N0M0<br>T3N0M0<br>T1N0M0<br>T2N0M0 | 1.14<br>112.92<br>0.81<br>22.36 | 0.175<br>48.2<br>0.533<br>5.014 | PR<br>PR<br>PR<br>PR | 2 / S |
| 7 | Mixed | NA | F | 1 | Abdominal | well differentiated | T2N1M0 | 104.6 | 104.6 | SD | 2 |
| 8 | Mixed | NA | F | 2 (a, b) | Abdominal (a)<br>Thorax lateral (b) | well differentiated<br>well differentiated | T3N0M0<br>T2N0M0 | 4.24<br>0.22 | 10.36<br>1.046 | PD<br>PD | 2 |
| 9 | English Pointers | 8 | F | 1 | Abdominal | well differentiated | T3N1M0 | 76.93 | 30.03 | PR | 2 / S |
| 10 | Boxer | 8 | F | 2 (a, b) | Thorax lateral (a)<br>Tibial (b) | well differentiated<br>well differentiated | T1N0M0<br>T1N0M0 | 0.314<br>0.141 | 0.113<br>0.28 | PR<br>PD | 2 |
| 11 | Mixed | 7 | F | 1 | Abdominal | well differentiated | T4N1M0 | 1.2 | 1.31 | SD | 1 |

M, male; F, female
PR: partial remission; SD: stable disease; PD: progressive disease
S: surgery
NR: death not related to cancer
NF: no follow-up
NA: not available

Based on the volume measurements, 11 (61.1%) lesions underwent PR, 4 (22.2%) underwent PD, and 3 (16.6%) underwent SD. We evaluated the tumor size at D0 as a prognostic factor according to the TNM recommendation for human SCCs, and we grouped the subjects according to whether they had tumors smaller or larger than 5 cm3. The volume at D0 had no impact on survival time (Fig 1B) and had no prognostic value (P>0.05).

In addition, the association between the clinical stage before treatment (D0) and the response to ECT was not significant (p=0.332) (Fig 1B). The median survival time of the 11 dogs was 180 days (32 to 570 days) (Fig 2A).

**Fig. 2.**
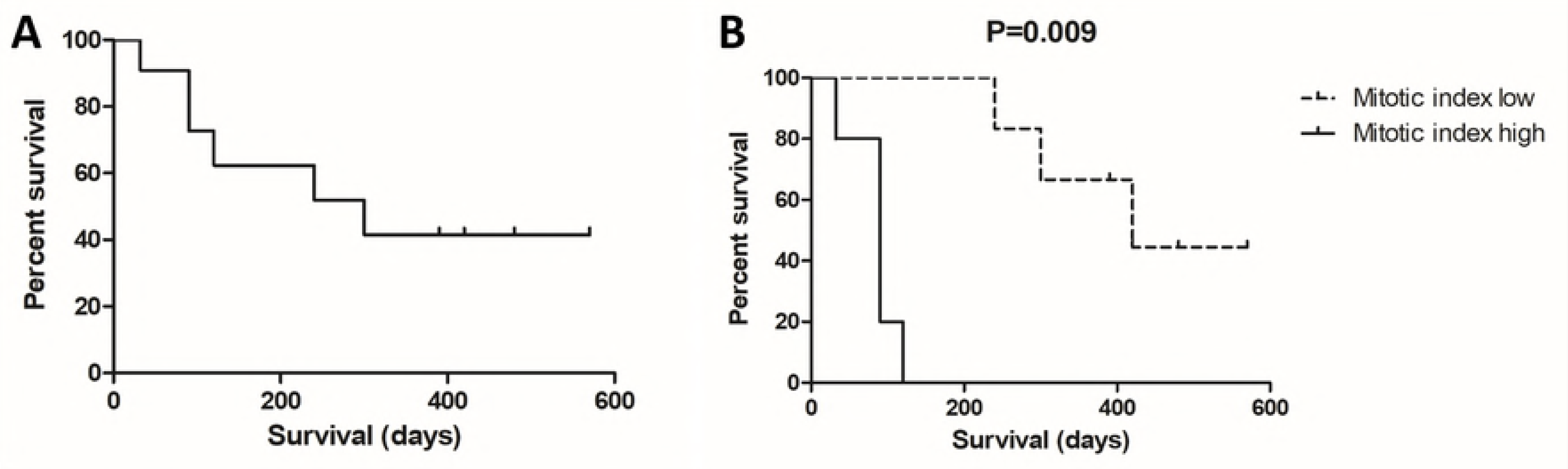
Overall median survival time of the 11 dogs and survival time based on mitotic index. (A) The median survival time of the 11 dogs was 180 days (32 to 570 days). (B) Subjects with mitotic index values lower than 4.9 at D0 experienced a longer survival time than subjects with mitotic index values greater than 4.9 (P=0.009).

The additional clinical results are shown in Table 1. The number of sessions that each subject underwent ranged from one to three. Five subjects underwent a surgical procedure after D21 (four due to PD and one due to the owner request).

### Histopathological features

Regarding the histopathological grade, 16 lesions (83.3%) were classified as well-differentiated SCC (Suppl Fig 2A), and three (16.6%) were classified as poorly differentiated SCC (Suppl Fig 2B). Interestingly, only well-differentiated tumors progressed, and all (3/18) poorly differentiated cSCC experienced PR.

The median mitotic index in the tumor group at D0 was 4.94 (0 to 34). We found a relationship between the mitotic index and overall survival. Subjects with mitosis numbers lower than 4.9 at D0 experienced a longer survival time (P=0.009) (Fig 2B).

Additionally, we found decreased mitotic index values at D21 compared to D0 (median 1.5, ranging from 0 to 20) (P=0.019) (Fig 1C). We also assessed intratumoral necrosis in the tumor samples at D0 and D21. It is not surprising that we found more necrosis in tumors at D21 than at D0 (P=0.041) (Fig 1D). We evaluated necrosis at D0 and the survival time, and we did not find significant differences (P>0.05).

### Protein expression

Immunohistochemical expression was evaluated in 15 tumors (15/18) at two different time points (D0 and D21). Three out of 18 samples were excluded due to the absence of neoplasticism in tumor samples from D0 or D21. The median expression of Ki67 in D0 was 277.96 (113.4 to 511.4), and D21 was 193.92 (15 to 494). We found decreased proliferative expression between D0 and D21 (Fig 3 A and B). Thus, tumor samples after ECT (D21) had a lower proliferative index than those before ECT (p=0.031) (Figure 1E). Using the proliferative index at D0, we evaluated the survival time between subjects with Ki67 values lower and higher than the median Ki67 value, and we did not find a significant difference (P>0.05).

**Figure 3.**
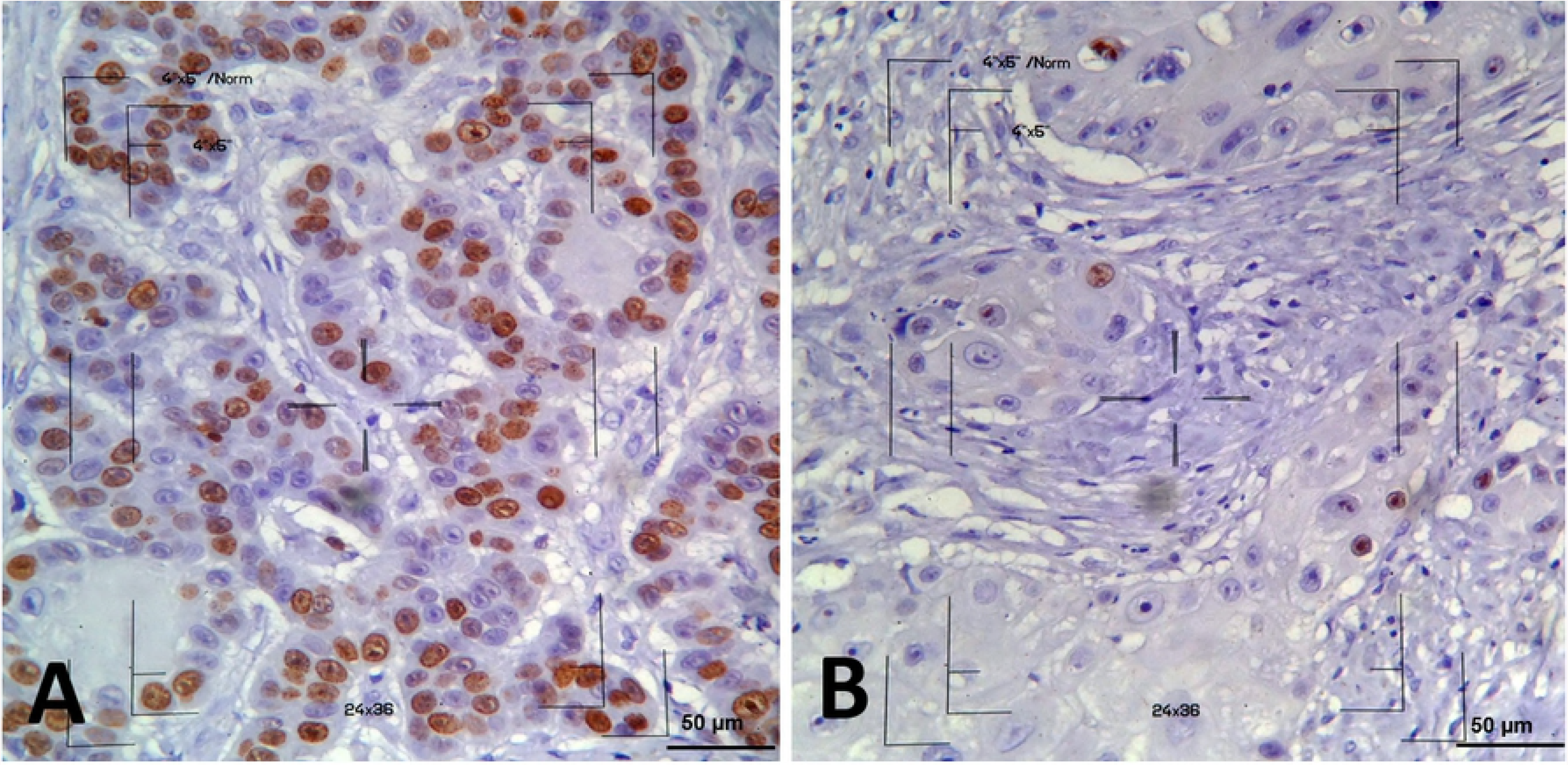
Photomicroscopy of cSCC in dogs with Ki67 immunolabeling. (A) High nuclear immunostaining of neoplastic cells at D0 (score 3). (B) Low nuclear immunostaining at D21 (score 1). The 4"x5" area, used for counting the cells, is observed in the images (immunocytochemistry, Envision, DAB, counterstaining with Harris hematoxylin, x400).

For BAX and Bcl-2, cytoplasmic expression was identified (Fig 4A). We found median BAX expression levels of 21.02% (ranging from 8.34 to 70.56) at D0 and 24.53% (ranging from 10 to 74.47) at D21. There was no significant difference in BAX expression between D0 and D21 (P>0.05) (Fig 4B). There was no significant difference in overall survival between subjects with low and high BAX expression levels (P>0.05) (Suppl Fig 1C).

**Fig. 4.**
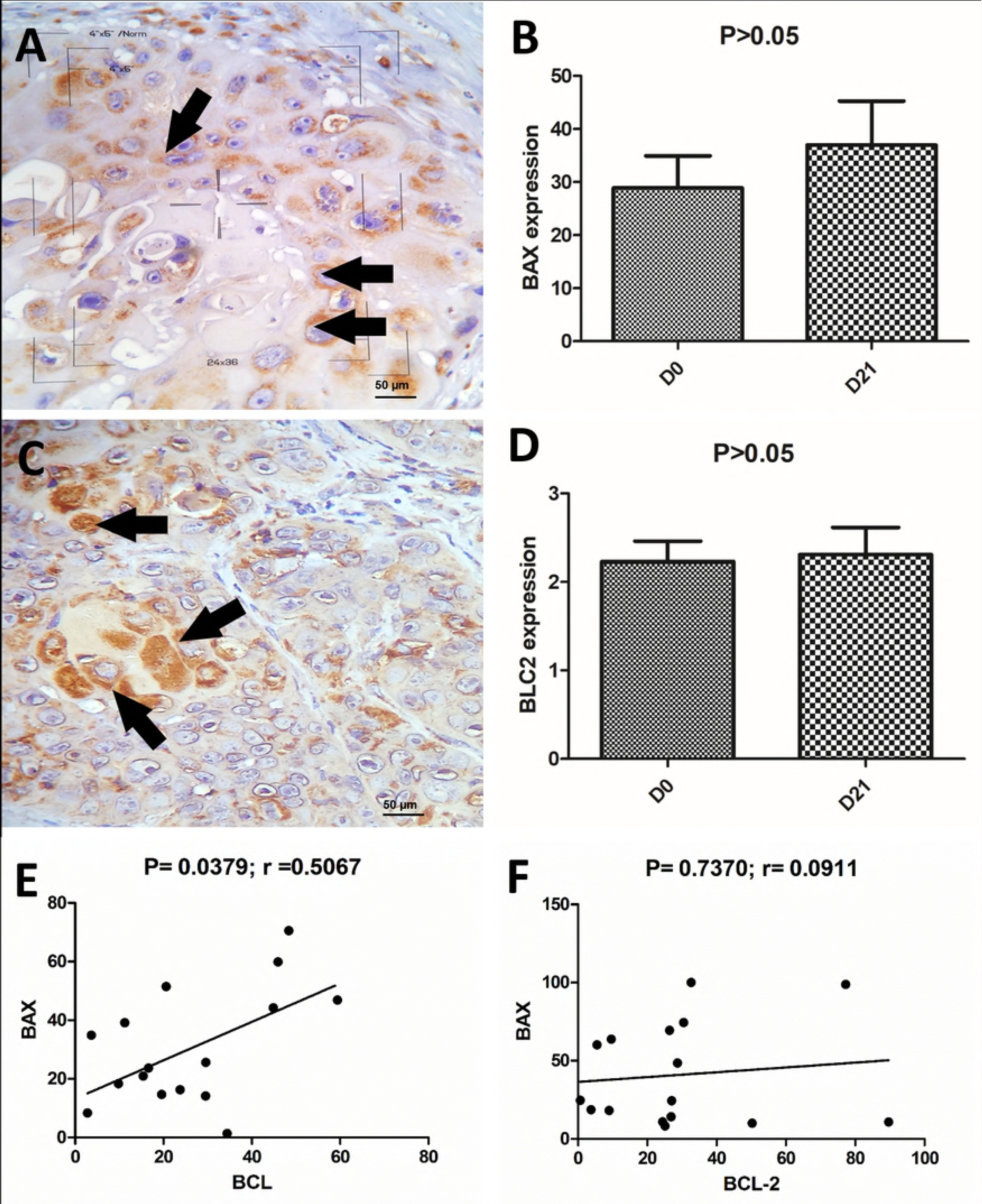
BAX and Bcl-2 immunoexpression levels and the correlation between both apoptotic markers in dogs with cSCC before and after ECT. (A) Cytoplasmic expression of BAX (arrows). (B) There was no significant difference in BAX expression between D0 and D21 (P>0.05). (C) Cytoplasmic expression of Bcl-2 (arrows). (D) There was no statistically significant difference in Bcl-2 expression between D0 and D21 (P>0.05). (E) Positive correlation between BAX and Bcl-2 before ECT (D0) (p=0.0379, r=0.5067). (F) Twenty-one days after treatment, no significant Gaussian approximation was observed (p=0.7370, r=0.0911)

**Fig. 5.**
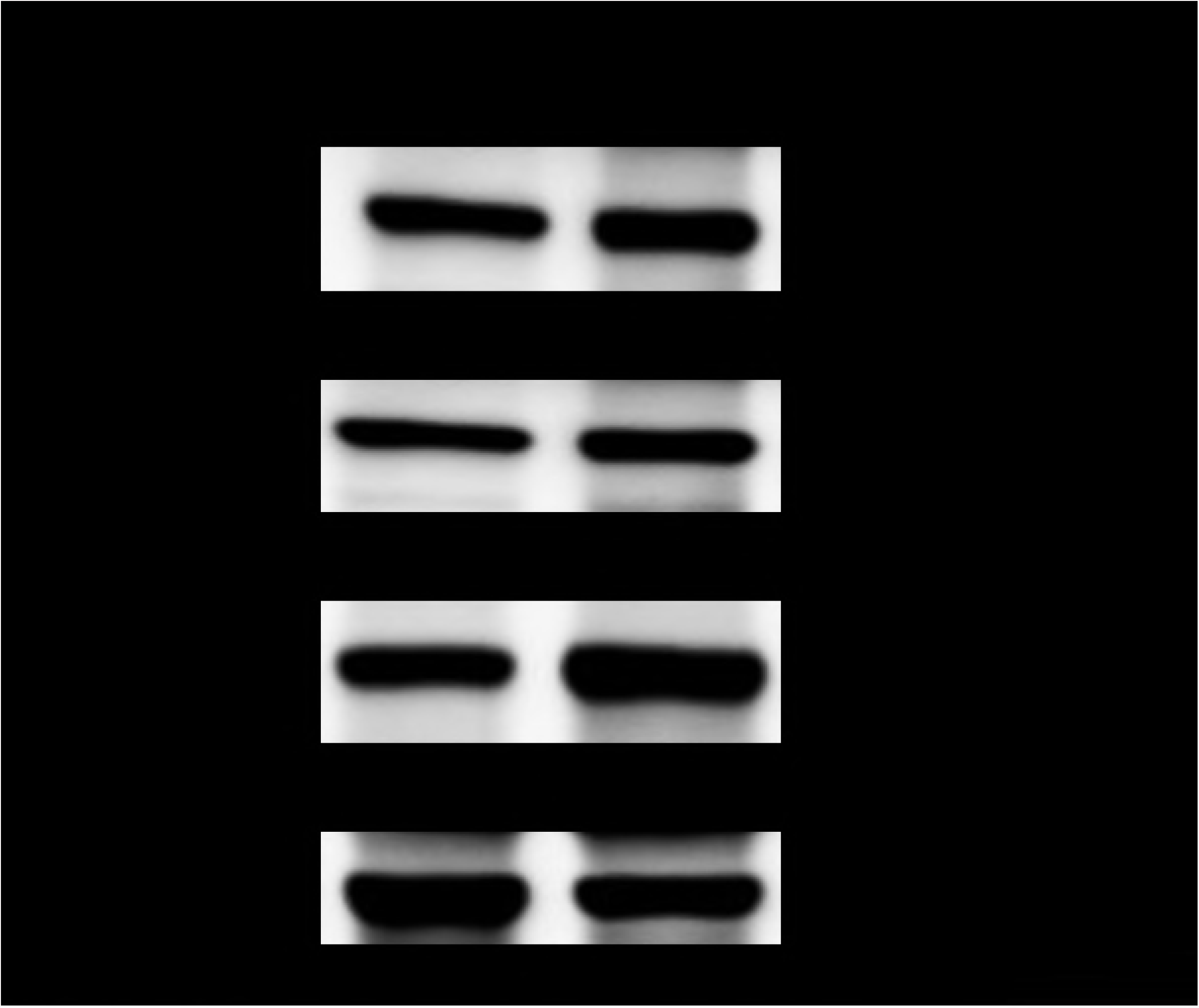

Regarding Bcl-2 expression (Fig 4C), we found medians of 20.62% (ranging from 2.76% to 48.37) at D0 and 26.42% at D21 (ranging from 0.65 to 77.23). There was no statistically significant difference in Bcl-2 expression between D0 and D21 (P>0.05) (Fig 4D). When we evaluated the survival time between samples with low Bcl-2 expression versus those with high expression, we did not find a significant difference (P>0.05) (Suppl Fig 1D).

Furthermore, a positive correlation was observed between BAX/Bcl-2 expression levels before ECT (D0) (p=0.0379, r=0.5067) (Fig 4E). However, after treatment (D21), we did not find a significant Gaussian approximation (p=0.7370, r=0.0911) (Fig 4F). Regarding the correlation between BAX/Ki67 and Bcl-2/Ki67, there was no significant difference among them at different time points (p>0.05).

### Western blot analysis

We found a 26-kDa band for Bcl-2 and a 21-kDa band size for BAX. Both antibodies showed only one band each in the western blot analysis (Fig 5).

## Discussion

cSCC is a very common tumor in tropical regions, and one of the biggest therapeutic challenges for tropical regions is removing the animal from exposure to sunlight during treatment. Because cSCC has a low level response to chemotherapy and a low metastatic rate, a local approach is required to achieve long-term disease-free interval and overall survival. Surgery is the primary treatment for cSCC. However, many patients have multiple tumors at the time of diagnosis, making surgical approaches difficult. Therefore, new techniques are required to improve the outcomes of canine patients with cSCC.

In this study, we evaluated ECT as a primary therapy for cSCC. We determined the association of different parameters with overall survival and tumor response to ECT. We demonstrated a reduction of the tumor size after ECT (D21), even in subjects with poorly differentiated tumors. Four lesions (in four different patients) presented progressive disease (PD). Of these four lesions, two had large volumes (ID 5 and 8), indicating that ECT may be a treatment option for small tumors. Unfortunately, we have no studies in Veterinary Medicine indicating the association of a tumor size cutoff with prognosis. In a large series study about the use of ECT in basal cell carcinoma (BCC) in humans, a 50% CR rate was observed after a single ECT cycle in primary BCC. Interestingly, in the same study, a second ECT cycle increased the CR rate from 50 to 63%, and retreatment was more advantageous in patients with local BCC. Thus, retreatment with ECT seems a reasonable option in patients with small BCCs to reduce the response duration [47]. In accordance with this human study, our results indicated that ECT could be a good therapeutic option for small tumors, and large may need more than one round of ECT. A previous case report by our study group observed complete remission of a digital trichoblastoma after three sessions of ECT, reaffirming the usefulness of repeated sessions of ECT for larger tumors [20].

Among the limiting factors of our study, there was an absence of a cutoff point for tumor size in cutaneous neoplasms in dogs, unlike in human medicine. In addition, the staging adopted in veterinary medicine does not take into account the size of the subject, which may influence the behavior of a tumor of the same size in dogs of different sizes. The small sample size is also a limitation of our study. However, we performed a G power analysis to guarantee that our tumor group had a sufficient number of samples.

Interestingly, lymph node involvement was not a prognostic factor in dogs with cSCC. The number and size of the lesions seems to be more important than the involvement of local lymph nodes. This can be explained by the invasive behavior of cSCC [5]. Usually, these tumors are more locally invasive than metastatic tumors [5,34,48]. On the other hand, the number of mitoses was correlated with overall survival in our study. Because the mitotic index is a routine evaluation, it will be important to evaluate the sensitivity and specificity of these indices to predict outcomes in dogs with cSCC. However, a large sample size is necessary. We also found more necrosis in samples after ECT, indicating that this therapy induces both apoptosis and necrosis. Thus, the greater amount of necrosis identified at D21 was due to the therapy.

One of the most interesting results of our research was the decreased proliferative index in tumors after ECT. Because ECT induces tumor apoptosis and necrosis, the number of proliferating cells was reduced. Furthermore, we did not find any difference in cell apoptosis between the two time points (D0 and D21), indicating that another mechanism may be involved in inhibiting tumor proliferation after ECT. Because ECT is a recent technique in Veterinary Medicine, a limited number of studies have demonstrated the ECT antitumoral effect in cSCC. In general, the high expression level of Ki67 observed in SCC, especially in poorly differentiated SCC, is related to an aggressive potential [9,34,37,49]. However, in the lesions studied, the proliferation index could also be considered high despite the predominance of well-differentiated SCC (83.3%), suggesting the aggressive potential, despite the degree of differentiation. Additionally, there is no established cutoff point for Ki67 to be considered an indicator of recurrence or metastasis, as occurs in cases of cutaneous mast cell tumor, which is 23 positive cells in a 10 mm x 10 mm / 400x area [50]. It should be noted that the complete evaluation of proliferation markers may be more effective in predicting the biological behavior of the tumor. In this context, Auler et al. [49] reported the case of a dog with well-differentiated SCC in the foreskin with inguinal lymph node metastasis. This animal showed high Ki67 positivity in the tumor tissue, in addition to the high expression of the growth factors HER-2 and EGFR (epidermal growth factor receptor).

A previous study observed the same pattern in mast cell tumors as the pattern identified in this study with regard to the proliferative index, which was observed in fewer positive cells for Ki67 among the neoplastic cells 28 days after treatment [19].

Regarding the apoptotic markers, all cSCC lesions showed cytoplasmic positivity for BAX. These results differ from those of Madewell et al. [28], who observed less than 2% BAX expression in cSCC in felines. However, other human studies have observed the expression of BAX in cutaneous and pulmonary SCC, which ranged from 53-100% [51,52]. A recent study observed that the expression of BAX in normal keratinocytes in the epidermis of dogs ranged from very weak to moderate, elucidating their importance in the pathogenesis of diseases [53]. To avoid any unspecific cross-reaction with canine tissue and the primary antibody, we performed western blotting for both antibodies using cSCC samples. Both antibodies presented only one specific band, confirming the reactivity of both antibodies with canine tissue.

Likewise, no significant difference was observed between the expression of the anti-apoptotic Bcl-2 protein before and after treatment with ECT. This is the first study to evaluate whether the immunohistochemical expression of Bcl-2 protein is altered by ECT in cSCC. A previous report evaluated the expression in mast cell tumors and observed increased expression of anti-Bcl-2 28 days after ECT and then decreased expression at D46 [19]. In our study, all cSCC samples showed immunolabeling of Bcl-2. However, other studies have shown different results and only one evaluated Bcl-1 by IHC in dogs with cSCC (n = 5), among which only one showed positivity [54]. In humans, Puizina-Ivic et al. [55] observed an absence of Bcl-2 positivity in SCC (n = 20), whereas Abu Juba et al. [56] observed positivity in 50% of cSCC samples (n = 10).

It must be noted that the correlation between BAX/Bcl-2 markers was significant before treatment, but no difference was observed afterwards, and no difference was observed between BAX/Ki67 and Bcl-2/Ki67 at both time points. In human studies, the Bax/Bcl-2 ratio can act as a rheostat that determines cell susceptibility to apoptosis [56], and lower levels of this ratio may lead to resistance of human cancer cells to apoptosis. In colorectal tumors, BAX and Bcl-2 expression levels were the most predictive of outcome when the Bax/Bcl-2 expression ratio was used [57]

Although the chemotherapeutic agents used have the capacity to cause cell damage that induces cell death by apoptosis, the BAX and Bcl-2 markers in the samples were not altered at the time point evaluated (D21). It is possible that other proteins involved in the apoptosis pathways exert a greater influence on the process of cell death induced by ECT. In addition, it cannot be ruled out that, in view of the dynamism and complexity of the apoptosis process, the expression of BAX and Bcl-2 may have been influenced by the ECT prior to the assessed moment, but not on the 21st day of treatment. A limitation in scoring a single protein is that the expression of a single protein may not reflect the level of apoptosis because apoptosis is a dynamic, complex process.

Regarding the proliferative index, 21 days after the application of ECT, it was possible to observe a significant reduction in proliferative indices in the lesions sampled. The same behavior was also observed in mammary neoplasms of human patients submitted to systemic chemotherapy because there was a significant reduction in the level of Ki67 in the tumor at 21 days of treatment [58,59]. However, in the veterinary literature, no research was found regarding the proliferative index before and after treatment in malignant neoplasms.

## Conclusions

ECT was able to reduce tumor volume and cellular proliferation in cSCC. Furthermore, our results showed that the BAX and Bcl-2 proteins did not show alterations in their expression levels in response to ECT at the time points evaluated. Therefore, studies involving the serial evaluation of these proteins, as well as the investigation of other proteins involved in apoptosis, may contribute to the understanding of the effect of ECT on apoptosis in canine cSCC.

---

**Suppl Fig. 1. Survival analysis according to lymph node involvement, tumor size, and apoptotic markers in dogs with cSCC.** (A) There was no difference in survival time between subjects with and without lymph node involvement (P > 0.05). (C) There was no significant difference in overall survival between subjects with high and low levels of BAX (P>0.05).

**Suppl Fig 2. Histopathological evaluation of squamous cell carcinoma (SCC) in dogs that underwent electrochemotherapy (ECT). (A) Well-differentiated SCC with the presence of mononuclear inflammatory infiltrate before ECT. (B) SCC before ECT. It is possible to see tissue disorganization with the presence of apoptotic cells showing basophilic nuclei.**

