## Supplementary figures and images for "Electrochemotherapy with bleomycin associated with doxorubicin induces tumor regression and decreases the proliferative index in canine cutaneous squamous cell carcinomas"

### Suppl Fig. 1

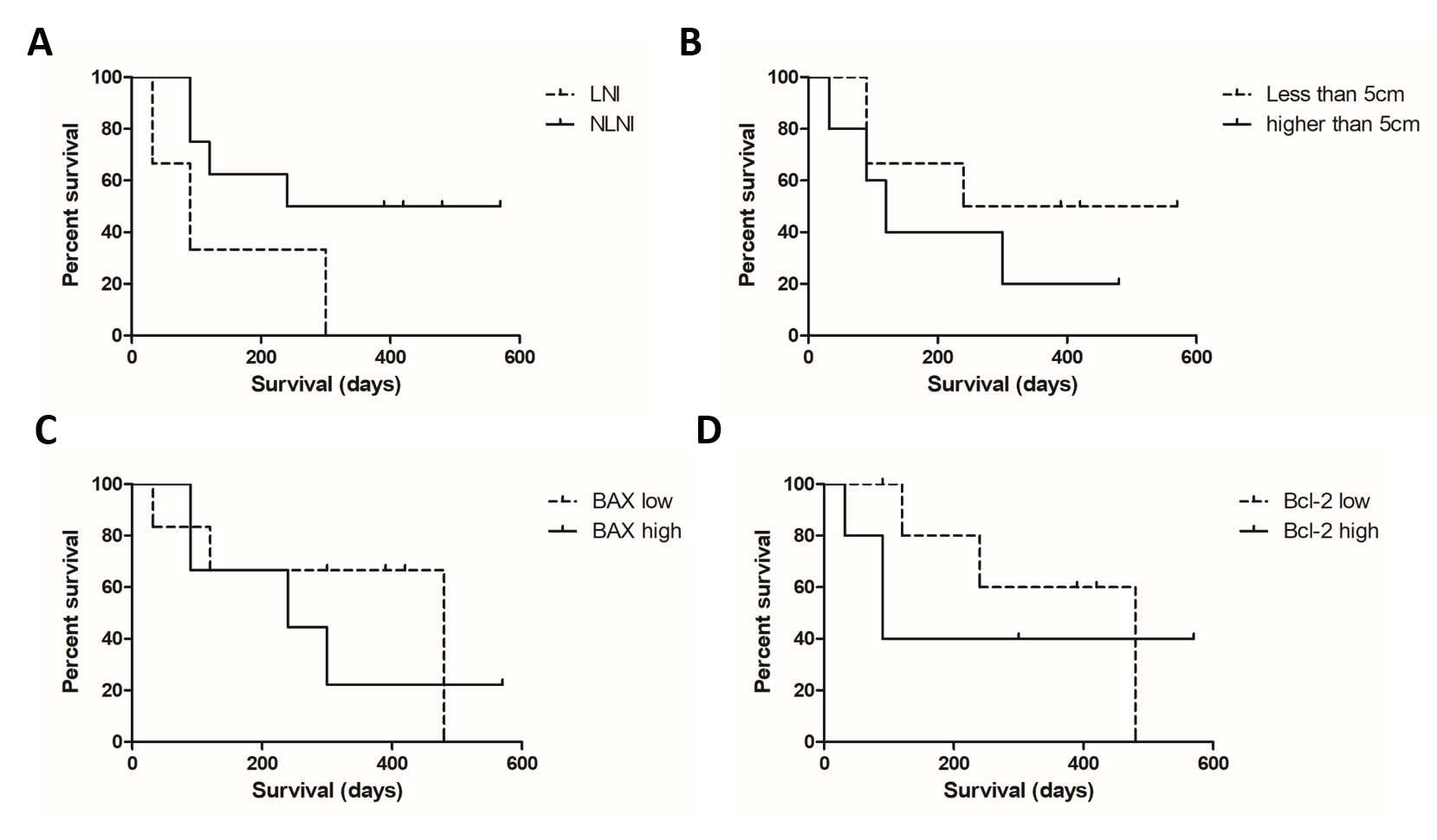

### Suppl Fig. 2

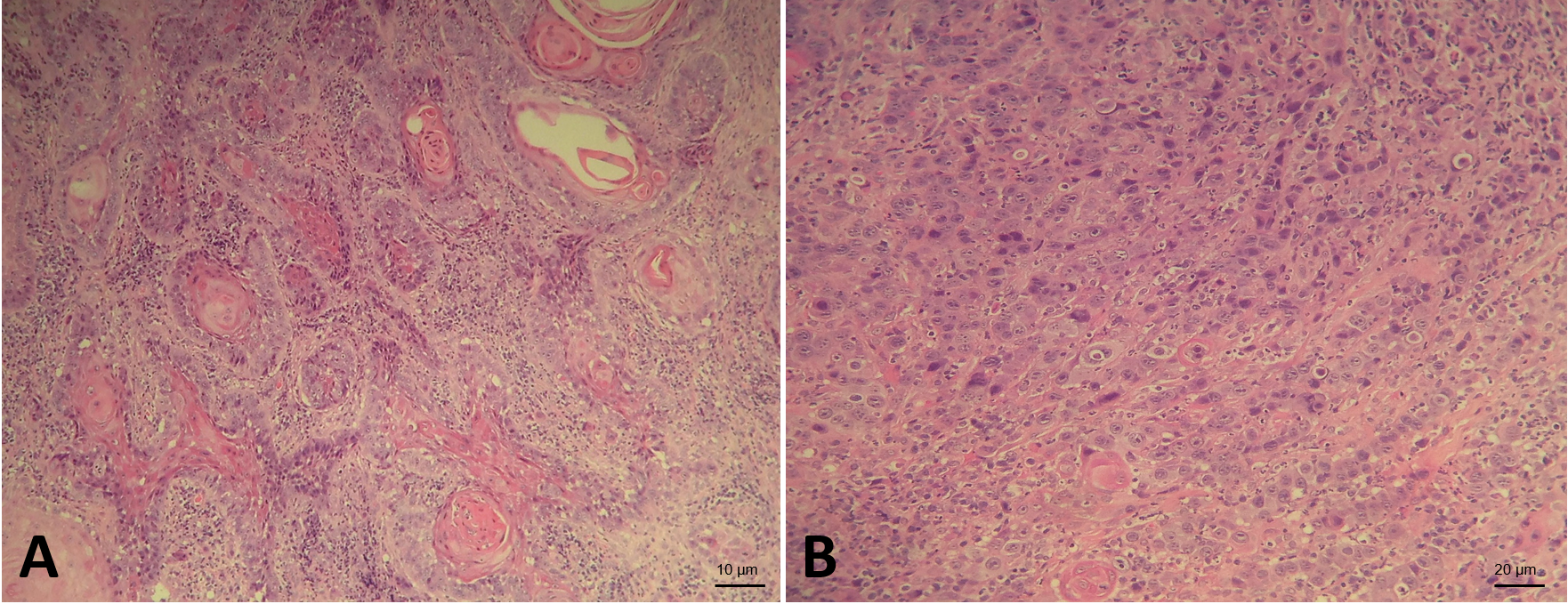
